# Targeted epigenomic changes to the maize methylome resulting from tissue culture

**DOI:** 10.1101/242081

**Authors:** Zhaoxue Han, Peter A. Crisp, Scott Stelpflug, Shawn M. Kaeppler, Qing Li, Nathan M. Springer

## Abstract

DNA methylation can contribute to the maintenance of genome integrity and regulation of gene expression. In most situations, DNA methylation patterns are inherited quite stably. However, changes in DNA methylation can occur at some loci as a result of tissue culture resulting in somaclonal variation. A sequence-capture bisulfite sequencing approach was implemented to monitor context-specific DNA methylation patterns in ~15Mb of the maize genome for a population of plants that had been regenerated from tissue culture. Plants that have been regenerated from tissue culture exhibit gains and losses of DNA methylation at a subset of genomic regions. There was evidence for a high rate of homozygous changes to DNA methylation levels that occur consistently in multiple independent tissue culture lines suggesting the existence of a targeted process for altering epigenetic state during tissue culture. The consistent changes induced by tissue culture include both gains and losses of DNA methylation and can affect CG, CHG or both contexts within a region. The majority of changes in DNA methylation exhibit stable inheritance although there is some evidence for stochastic reacquisition of the initial epigenetic state in some individuals. This study provides insights into the susceptibility of some loci and potential mechanisms that could contribute to altered DNA methylation and epigenetic state that occur during tissue culture in plant species.

## Introduction

Tissue culture is used in many commercially important plant species, both for clonal propagation as well as successful transformation and regeneration of transgenic materials. While it is expected that clonal plants derived from tissue culture will have no changes in genetic information there are frequent examples of somaclonal variation, manifested as heritable phenotypic differences in plants recovered from tissue culture (Phillips *et al*. 1994; Kaeppler *et al*. 2000; Miguel and Marum 2011; Neelakandan and Wang 2012). In some cases, this could be due to activation of transposons during tissue culture (Peschke *et al*. 1987; Hirochika 1993). However, in other cases there is evidence for epigenetic changes that are directly linked to gene expression variation manifested as phenotypic variation in plants derived from culture (Rhee *et al*. 2010; Ong-Abdullah *et al*. 2015).

In plants, DNA methylation variation can be associated with heritable differences in gene expression in the absence of DNA sequence changes. Approximately 30% of the cytosines in the maize genome are present as 5-methylcytosine (Papa *et al*. 2001). DNA methylation is particularly prevalent in CG dinucleotide or CHG (where H is A, C or T) trinucleotide contexts, with lower levels of DNA methylation at CHH sites (Gent *et al*. 2013; Regulski *et al*. 2013; West *et al*. 2014). There are distinct mechanisms to target and maintain methylation in each of these sequence contexts (Law and Jacobsen 2010; Matzke and Mosher 2014; Springer and Schmitz 2017). Maintenance pathways are required to re-methylate daughter strands following DNA replication otherwise DNA methylation can be passively lost over rounds of replication. Plants also utilize active mechanisms for demethylation (Zhang and Zhu 2012) and the plant methylome is the result of combined methylation and demethylation activities.

The sources and prevalence of epigenetic variation are of significant interest. There is abundant evidence for natural variation in DNA methylation in plants, as shown in in maize and other plant species (Taudt *et al*. 2016; Springer and Schmitz 2017). There is less evidence for developmental variation in DNA methylation for most vegetative tissues (Eichten SR 2013; Schmitz *et al*. 2013; Li *et al*. 2015c; Kawakatsu *et al*. 2016). While several studies have provided examples of DNA methylation changes induced by the environment (Dowen *et al*. 2012; Jiang *et al*. 2014; Wibowo *et al*. 2016) other studies have found quite limited evidence for changes in DNA methylation in response to the environment (Song *et al*. 2013; Hagmann *et al*. 2015; Eichten and Springer 2015; Secco *et al*. 2015; Crisp *et al*. 2017; Ganguly *et al*. 2017).

There is evidence that DNA methylation patterns are perturbed by tissue culture. Previous studies have found locus-specific effects on DNA methylation (Kaeppler and Phillips 1993) and several epialleles resulting from tissue culture have been characterized (Krizova *et al*. 2009; Rhee *et al*. 2010; Ong-Abdullah *et al*. 2015). Recent studies have provided genome-wide evidence for changes in DNA methylation following tissue culture in Arabidopsis, rice and maize (Tanurdzic *et al*. 2008; Stroud *et al*. 2013; Stelpflug *et al*. 2014). Each of these studies have documented hundreds of loci with altered DNA methylation levels following tissue culture and have found evidence for losses of DNA methylation that are consistent in independent tissue culture derived lines. Some epialleles can generate new phenotypes, for instance the *Karma* epiallele in oil palm is associated with loss of CHG methylation at a transposable element (TE) nested in a gene (Ong-Abdullah *et al*. 2015) resulting in production of aberrant transcripts. There is also evidence for Mendelian inheritance of epialleles following tissue culture in rice and maize (Stroud *et al*. 2013; Stelpflug *et al*. 2014). However, the mechanistic basis for the observed perturbations in DNA methylation remain largely unknown.

Previous work on maize utilized a quantitative approach to document locus-specific methylation levels by coupling immunoprecipitation of methylated DNA with hybridization to microarrays (Stelpflug *et al*. 2014). While this method can be useful for documenting regions with quantitative variation in DNA methylation, the procedure lacks the ability to resolve context-specific differences and lacks the resolution for determining the specific boundaries of altered methylation. In this study, we utilized a sequence-capture bisulfite sequencing approach to profile context specific DNA methylation levels in 25 R_1_ plants recovered from tissue culture. We find many regions with altered DNA methylation levels, particularly in the CG and CHG context. A subset of the changes in DNA methylation are consistently found in many independent events and appear to represent homozygous changes in DNA methylation that occur during tissue culture. The majority of these consistent changes in methylation are also observed to occur in natural populations. Analysis of context-specific changes in methylation allowed us to identify numerous CG-only and CHG-only differentially methylated regions (DMRs). Moreover, we document multiple examples of Karma-like (Ong-Abdullah *et al*. 2015) epialleles with context-specific changes in DNA methylation. At these loci, heterochromatic DNA methylation is lost at transposable elements located within genes. This study may shed light on the mechanisms that drive epigenome changes during tissue culture.

## Results

We employed a strategy that coupled bisulfite modification with sequence capture termed “SeqCap-Epi-v2” to document context-specific DNA methylation patterns of the maize genome (AGPv4 - (Jiao *et al*. 2017)). The design captured 22,749 specific target regions covering 15.7 Mb (see https://doi.org/10.13020/D69X0H and methods for detailed description of the sequence capture design). Methylation patterns were profiled in three sibling A188 plants that had not passed through tissue culture as well as one callus sample, one primary R_0_ regenerant and 25 R_1_ plants that resulted from self-pollination of primary regenerants (**Figure 1; Table S1**). The level of DNA methylation in the CG, CHG and CHH context was determined for each region in each sample and we focused our analysis on 15,325 regions with coverage of at least 3 reads and consistent methylation levels among the three control siblings (see methods). Regions with >30% variation between control samples in the CG or CHG context, or >10% in the CHH context, were excluded. We reasoned that these regions likely represent metastable loci unrelated to tissue culture treatment but may be of interest in future analysis. In addition, we profiled five inbreds from diverse backgrounds for comparison including three biological replicates of B73, 2 replicates of Mo17, and single replicates of W22, MoG and Ki11 (**Table S1**).

**Figure 1.**
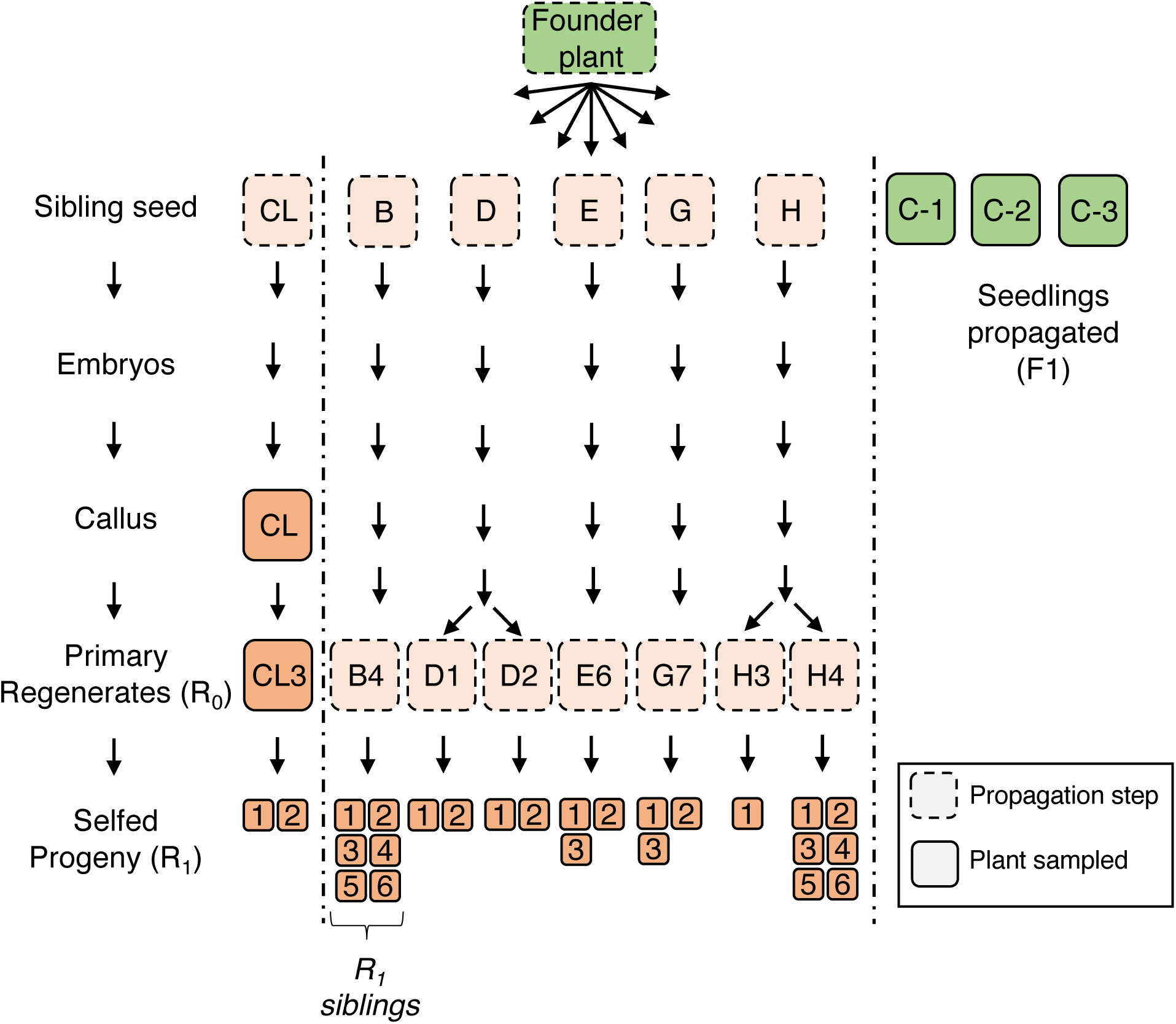
Transgenerational tissue culture experimental design. Overview of the tissue-culture experimental design comprised of six cell-lines initiated from independent embryos originating from the same progenitor plant. Samples are represented by the squares, solid outlines indicate samples harvested for DNA methylation profiling.

### Frequent hypomethylation in CG and CHG contexts following tissue culture

Genome-wide methylation patterns were assessed in A188 plants before, during and after tissue culture, and were subsequently contrasted with five maize inbreds from diverse genetic backgrounds. Principal component analysis of CG methylation revealed clustering driven largely by genotype (**Figure 2A**). The tissue culture samples were generally clustered closely with A188 plants that had not experienced tissue culture. However, PC2 does differentiate plants that have been passed through tissue culture from those that have not (**Figure 2A**). Differentially methylated regions (DMRs) were identified for thorough analysis of context-specific DNA methylation levels for each region in each sample relative to the controls (**Figure 2B, Figure S1**). For CG and CHG methylation, DMRs are defined as regions where the DNA methylation percentage levels differ by greater than 40% in a sample relative to the control average. DMRs in the CHH context were identified when one sample has >25% and another has less than 5% CHH methylation. In total 3,921 distinct DMRs were identified. There are many more CG and CHG DMRs than CHH DMRs even though less strict criteria are used for CHH DMR discovery (**Figure 2B, Figure S1**). CHH DMRs are more likely to represent hypermethylation if a sample that has experienced tissue culture relative to the controls, while CG and CHG DMRs are more likely to exhibit hypomethylation in the tissue culture samples (**Figure 2B**).

**Figure 2.**
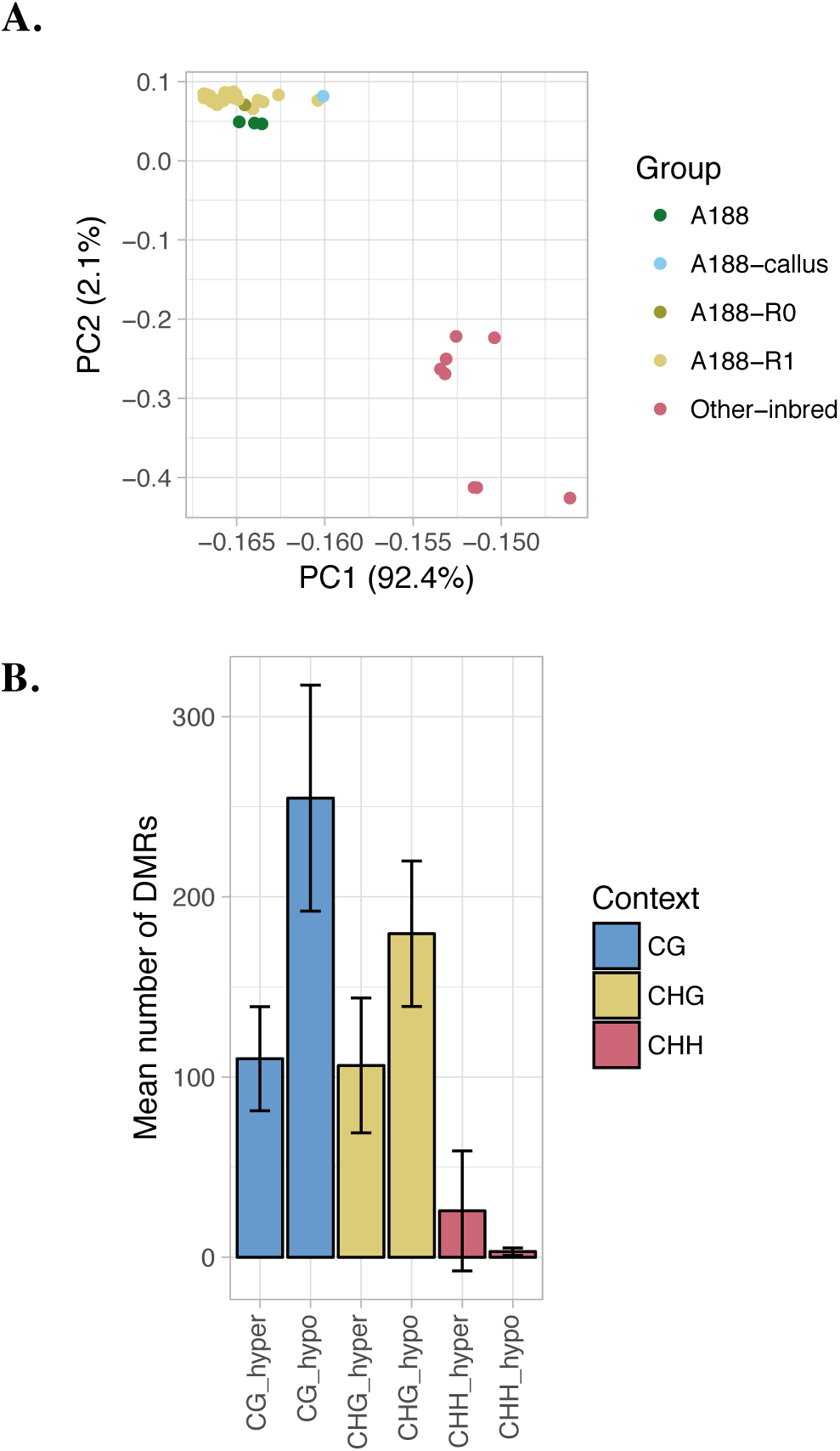
Frequent hypomethylation in CG and CHG contexts following tissue culture. (A) Principal component analysis of CG methylation levels for A188 samples from the tissue culture experiment (non-cultured seedling, callus, R_0_ and R_1_ regenerated seedlings) contrasted with other inbreds (B73, Mo17, MoG, Ki11 and W22). (B) The average number of DMRs per context identified for samples passed through tissue culture. Bars indicate the average number of DMRs for both hypermethylated and hypomethylated regions. The error bars report the standard deviation for DMR numbers across all samples.

For all DMRs the relative levels of DNA methylation in each context were examined to identify the relative frequency of regions where DNA methylation changes in multiple contexts as opposed to changes that are context-specific (**Figure 3; Figure S2**). There are very few examples of changes in CHH methylation levels at CG or CHG DMRs and in most cases the level of CHH methylation at these regions is very low in both the control and tissue culture samples (**Figure S2**). There is also limited evidence for changes in CG or CHG methylation at the majority of CHH DMRs (**Figure S2**). Therefore, CHH DMRs seem to represent a unique set of genomic regions that have limited overlap with changes in methylation in other contexts. In contrast, many of the CG or CHG DMRs also exhibit similar changes in the other context (**Figure 3A; Figure S2A**). Each CG/CHG DMR was classified as CG-only, CHG-only, or CG-CHG based on the relative levels of DNA methylation in each context (summarized in **Table S2**). This classification was limited to regions for which there was data in both contexts to enable determination of context specificity. In total, 1,255 distinct regions out of 2939 total CG and CHG DMRs were characterized as CG-only, CHG-only, or dual CG-CHG DMRs in this manner.

**Figure 3.**
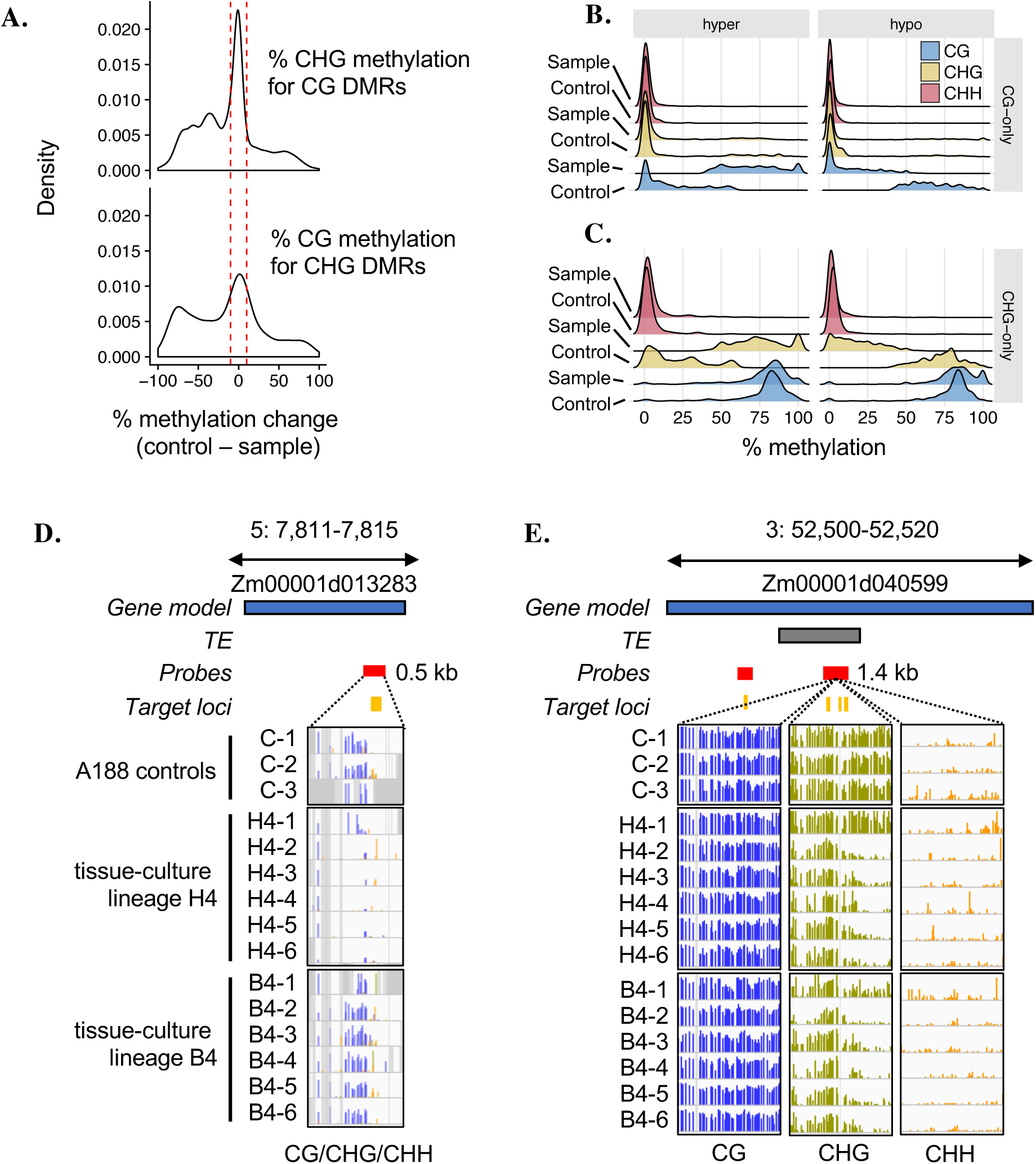
Identification of context specific DMRs resulting from tissue culture. (A) Density plots of methylation changes (sample – control) for tissue culture sample B4_1 for regions identified as DMRs in a particular context. This plot illustrates that CG and CHG levels exhibit two behaviours: they can change independently (context specific DMRs) as indicated by the areas between the red lines; and also concurrently. (B)-(C) Overlayed density plots (ribbon plots) summarising the distributions of methylation levels in each context at all CG-only and CHG-only DMRs identified in regenerated plants. The y axis represents arbitrary density units. (D) Example of a gene body CG-only DMR in the B4 and H4 tissue-culture lineages. The blue and grey boxes represent the relative positions of gene and TE models (v4 genome annotation); the red and yellow boxes indicate the SeqCap-Epi_v2 capture probes locations and target loci respectively. For each sample track, bar height represents % methylation (0–100%), purple = CG, green = CHG and yellow = CHH. Grey shading indicates missing data not captured. (E) Example of a *karma*-like CHG-only hypomethylation at a TE nested in gene Zm00001d040599.

The maize B73 genome can be broadly divided into four main methylation domains, characterized by distinct methylation profiles (noting that more sophisticated definitions have also been proposed (Gouil and Baulcombe 2016)). A small fraction (3.9%) is unmethylated in all contexts; 0.7% has high CHH and represents RdDM targets, 7.1% has high CG-only and is found in gene bodies, and the largest fraction (47.9%) has high levels of both CG and CHG and represents heterochromatin (Springer and Schmitz 2017). The remaining 40.3% of the maize genome is unclassified due to either a lack of coverage or intermediate methylation levels. The SeqCap-Epi-v2 design does not uniformly target these regions and the captured portions of the genome include 8.3% RdDM targets, 12.1% high CG-only, 50.6% heterochromatin, 11.6% unmethylated and 10.4% unknown.

We assessed how the changes in methylation induced by tissue culture affected the shifts among these types of domains. For instance, the CG-CHG DMRs largely represented a transformation of heterochromatic region in control plants to unmethylated domains in regenerated plants. Further examination of DNA methylation in both CG and CHG contexts for the CG-only or CHG-only DMRs revealed interesting patterns (**Figure 3B-C**). The CG-only DMRs - irrespective of whether they were hyper- or hypo-methylated in the tissue culture samples - were predominantly unmethylated in the CHG context in both the control and tissue culture sample, characteristic of the gene-body methylation domain (**Figure 3B)**. These CG-only DMRs therefore reflect shifts between the unmethylated and CG-only methylation. In contrast, CHG-only DMRs that exhibit either hyper- or hypo-methylation in the tissue culture sample were frequently methylated in the CG context in both samples, characteristic of heterochromatin (**Figure 3C**). These CHG-only DMRs represent interconversion between heterochromatin and “gene body-like” CG-only methylation.

Examples for a CG-only and a CHG-only DMR are shown in **Figure 3D-E**. The CG-only DMR in **Figure 3D** occurs within a gene body and represents a loss of CG methylation. This loss is seen in five of the six H4-family siblings but one individual has partial CG methylation, likely indicating that it has regained methylation. None of the individuals in the B4 lineage exhibit changes in CG methylation at this locus. The CHG-only DMR in **Figure 3E** occurs in a region that has high levels of CG and CHG methylation in the control plants. CHG methylation is substantially reduced, especially at the 3’ region, in all the B4 individuals and in five of the six H4 individuals. However, one individual (H4-1) appears to have largely regained CHG methylation. Interestingly, this CHG DMR is flanked by CHG methylation that is retained. It is possible that this methylation can provide a “seed” for the return of methylation through spreading. These patterns suggest the potential for consistent changes during culture and a stochastic reacquisition of DNA methylation in some individuals similar to the slow reacquisition of methylation at some DMRs in the epiRIL population (Catoni *et al*. 2017).

### A subset of methylation changes induced by tissue culture are common to many independent samples

A non-redundant set of DMRs that were identified in at least one tissue culture sample were further assessed to document the frequency of shared changes in methylation in multiple related, or unrelated, plants that had passed through tissue culture (**Figure 4**). On average, there were approximately 500 DMRs per sample out of 15,325 regions profiled (3.3% on average). The probability of one DMR occurring in 14 or more (>50%) samples is 5.1 × 10^−20^. Thus, in the absence of a targeting mechanism directing heritable changes in DNA methylation the null expectation is that DMRs would occur randomly in the genome with virtually no conservation between samples. However, we observe conservation of the same DMRs in up to 27 samples. The number of recurring DMRs was observed at a frequency far exceeding the level of overlap predicted by simulations (**Figure S3A**). For instance, given the total number of observed CG-hypo DMRs per sample, we would expect by random chance to find less than one DMR to recur in 6 or more samples (p << 0.001), yet we observe 303 DMRs common to six or more samples.

**Figure 4.**
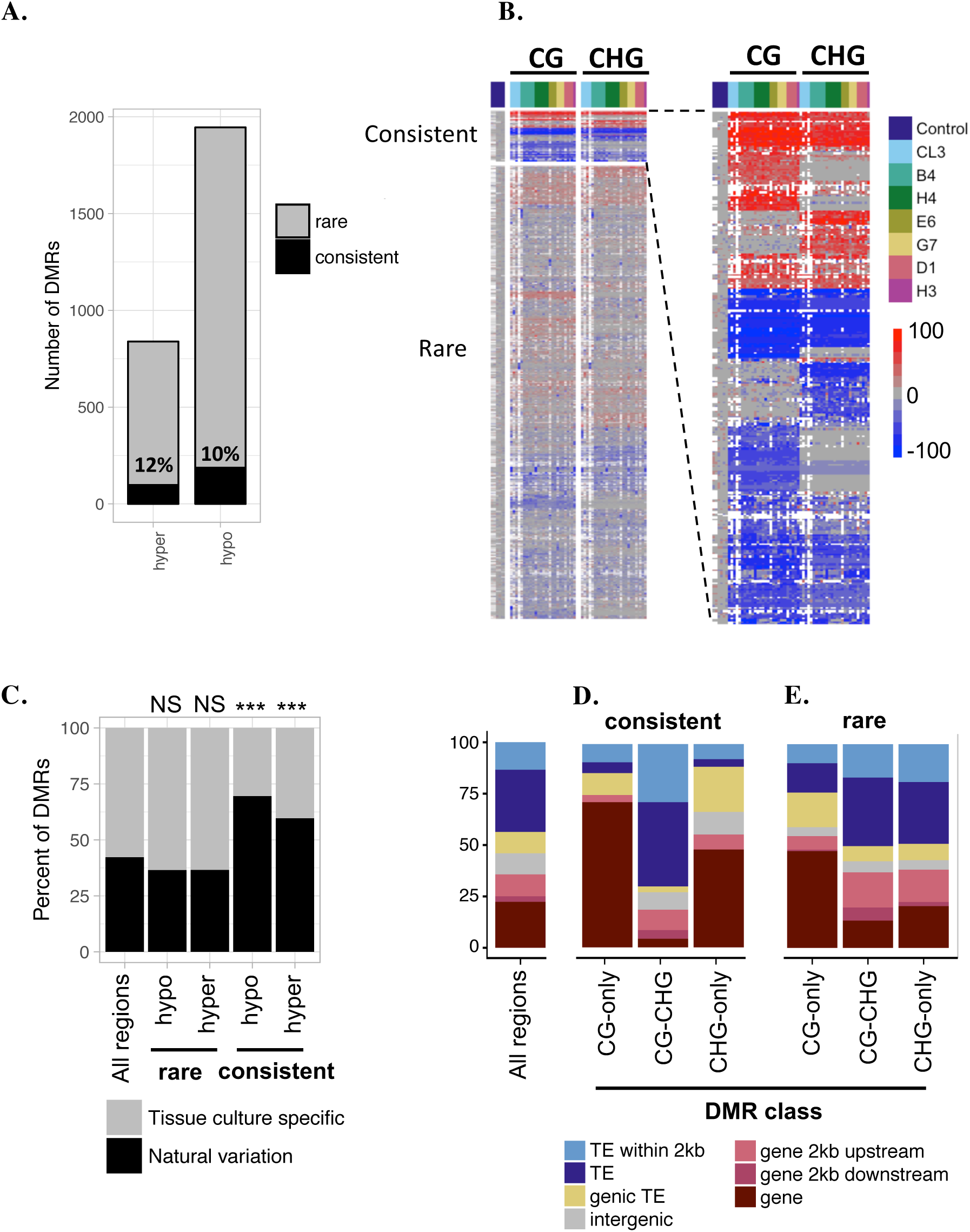
A subset of methylation changes induced by tissue culture are common to many Independent samples (A) The relative numbers of rare and consistent DMRs combined for the CG and CHG context are shown. The direction of a DMR (hypo/hyper) was assigned If there was >80% agreement between samples, and the loci was considered consistent If >50% of samples exhibited the DMR. Numbers Indicate the percentage of consistent DMRs In each category. (B) Hierarchical clustering of the change In methylation levels (sample-control) for all samples at each DMR In the CG and CHG context. White shading denotes missing data. (C) The percentage of tissue culture DMRs that are observed to occur naturally among a panel of 19 Inbreds Is shown (*** p < 0.001; NS, not significant, hypergeometrlc test). (D)-(E) The genomic feature(s) overlapping each DMR – categorized as (D) consistent or (E) rare – were determined and the percentage of each feature Is shown. The distribution of the background s*et al*l regions Is shown to the left of each panel.

DMRs that were identified in <50% of samples were classified as “rare” while DMRs identified in >50% of samples were classified as “consistent” (**Table 1 and Table S3**). The majority of rare DMRs are only observed in one or two samples while there are a set of the consistent DMRs found in nearly all samples (**Figure S3B**). As expected, CHH DMRs were almost entirely rare (99%). Nevertheless, 5 out of 424 CHH DMRs were found to be consistent, which is more than expected by chance alone. Given the number of CHH DMRs, we would not expect to find any DMRs recurring in more than 2 samples (p << 0.001). A much larger subset of CG and CHG DMRs display strong consistency between samples. Here rare and consistent DMRs occur in the same proportions for hypo and hyper directionality, with rare comprising around 90% and consistent 10% of total DMRs identified (**Figure 4A**). The distinction between rare and consistent DMRs can also be observed when methylation levels for all samples are hierarchically clustered (**Figure 4B**). In each individual sample approximately half the DMRs identified are consistent and shared with other samples, while half are rare (**Figure S3C**). An example of the DNA methylation patterns exhibited in a common DMR is shown in **Figure S3D**. In some cases, DMRs classified as rare occur where only a few samples reach the significance threshold with the remaining samples demonstrating consistent but partial loss or gain of methylation (**Figure 4B**), reflecting the relatively high stringency of our DMR identification. Thus, the DMRs identified as consistent represent a high confidence set of recurring DMRs.

**Table 1.** Characterization of CG, CHG and CHH DMRs

| Specificity consensus | Direction consensus | Consistency | Number of regions | % natural variants |
| --- | --- | --- | --- | --- |
| CG-only | hyper | consistent | 21 | 76.1 |
|  |  | rare | 152 | 59.6 |
|  | hypo | consistent | 36 | 85.7 |
|  |  | rare | 380 | 52.3 |
| CG-CHG | hyper | consistent | 22 | 54.5 |
|  |  | rare | 37 | 51.4 |
|  | hypo | consistent | 49 | 79.1 |
|  |  | rare | 165 | 34.5 |
| CHG-only | hyper | consistent | 12 | 100 |
|  |  | rare | 114 | 40.7 |
|  | hypo | consistent | 16 | 81.3 |
|  |  | rare | 236 | 40.3 |
| unclassified | hyper | consistent | 43 | 48.3 |
|  |  | rare | 438 | 30.0 |
|  | hypo | consistent | 86 | 59.3 |
|  |  | rare | 977 | 31.9 |
| CHH | hyper | consistent | 4 | 1.1 |
|  |  | rare | 372 | 98.9 |
|  | hypo | consistent | 1 | 2.1 |
|  |  | rare | 47 | 97.9 |

All CG, CHG and CHH DMRs were categorized based on the methylation context consensus and consistency between samples. CHH DMRs were analyzed separately from CG and CHG. For CG and CHG specificity and consistency a consensus of $\geq 80\%$ of samples was required or else regions were denoted “unclassified”. If there was $\geq 50\%$ support for a DMR between samples it was classified as “consistent”, all others were considered “rare”.

### Features of consistent DMRs

The features of CG and CHG DMRs were further documented to better understand why some regions exhibit consistent changes in methylation given that many other DMRs behave more stochastically. The tissue culture induced DMRs were compared to a set of naturally occurring DMRs found among a wider panel of 19 inbreds subject to whole genome bisulfite sequencing (WGBS) (Anderson *et al*. 2017). Consistent DMRs exhibited a significant overlap with loci that exhibit natural variation (**Figure 4C** p < 0.001, hypergeometric test). In contrast, the rare DMRs do not exhibit significant enrichment or depletion for overlap with natural variation DMRs (p > 0.5, hypergeometric test). For example, the DMR in **Figure S3D** shows an region with low methylation in A188 and B73 but high levels of DNA methylation in the other inbreds and in all samples that have been passed through tissue culture. The concurrence of these A188 tissue culture DMRs with natural variants across a wide panel of inbreds, possibly suggests that these DMRs represent examples of pure epigenetic variation in which DNA methylation variation is not due to genetic changes among inbred lines of maize.

Next we examined the genomic locations of consistent CG and CHG DMRs. For instance, given that CG-only DMRs occur where CHG is low, we hypothesized that these DMRs would be enriched in gene bodies, and given that CHG-only DMRs occur where CG is high (characteristic of heterochromatin) we anticipated enrichment at TEs and intergenic regions. Consistent DMRs displayed significant bias in their location in the genome (**Figure 4D**). Consistent CG-only and CHG-only DMRs were both enriched in genes and depleted in TEs (FDR < 0.05, hypergeometric test), while consistent CG-CHG DMRs were depleted at genes (FDR < 0.05) and enriched at TEs (although not statistically significant after FDR correction). Rare CG-only DMRs had a similar profile to consistent CG-only DMRs. By contrast, rare CG-CHG/CHG-only DMRs had a profile more consistent with the background frequency, with some enrichment in promoters (FDR < 0.05) and close to TEs (FDR < 0.05; **Figure 4E**).

The enrichment of CG-only DMRs in genes was consistent with our expectation that these DMRs have the characteristics of gene-body methylation. It was somewhat unexpected to also find CHG-only DMRs enriched in genes given that these DMRs occur in a methylation context characteristic of heterochromatin. In total, 64% (18/28) of the consistent CHG-only DMRs occur at locations of genic heterochromatin. We also observed that many of these genes have nested TEs (for example **Figure 3E and Figure 5F**) reminiscent of the *Karma* locus in oil palm (Ong-Abdullah *et al*. 2015). This prompted us to examine the prevalence of potential *Karma-*like epialleles in more detail. Indeed, the example shown in **Figure 3E** of a CHG-only DMR occurs within a TE that is located within a gene. Across the maize genome over 10% of maize genes contain at least one TE insertion over 1 kb within an intron (West *et al*. 2014; Hirsch *et al*. 2016). In total, our sequence capture design profiled methylation information at 1,447 genic loci that have high CG and CHG methylation in the control A188 plants indicative of heterochromatin (control average >50% CG and >50% CHG). We detected significantly reduced levels of CHG methylation at 200 of these loci (219 DMRs, **Table S4**). These genes were found to have diverse functions, and Gene Ontology (GO) analysis revealed that several genes were linked to nucleotide metabolic processes (eg, GO:0009165 and GO:0055086); although, following FDR correction, no GO categories were statistically enriched relative to the background set of genes profiled. The heterochromatic DMRs were mostly rare (187/219); nevertheless, 37 (20%) were consistently observed in multiple independent plants (**Table S5**). Notably, the consistent set included genes involved in developmental processes that could possibly impact somatic embryogenesis including, Histone H2A (Zm00001d012837), *meg4* (Zm00001d019541) and *Hox1a* (Zm00001d010758)

**Figure 5.**
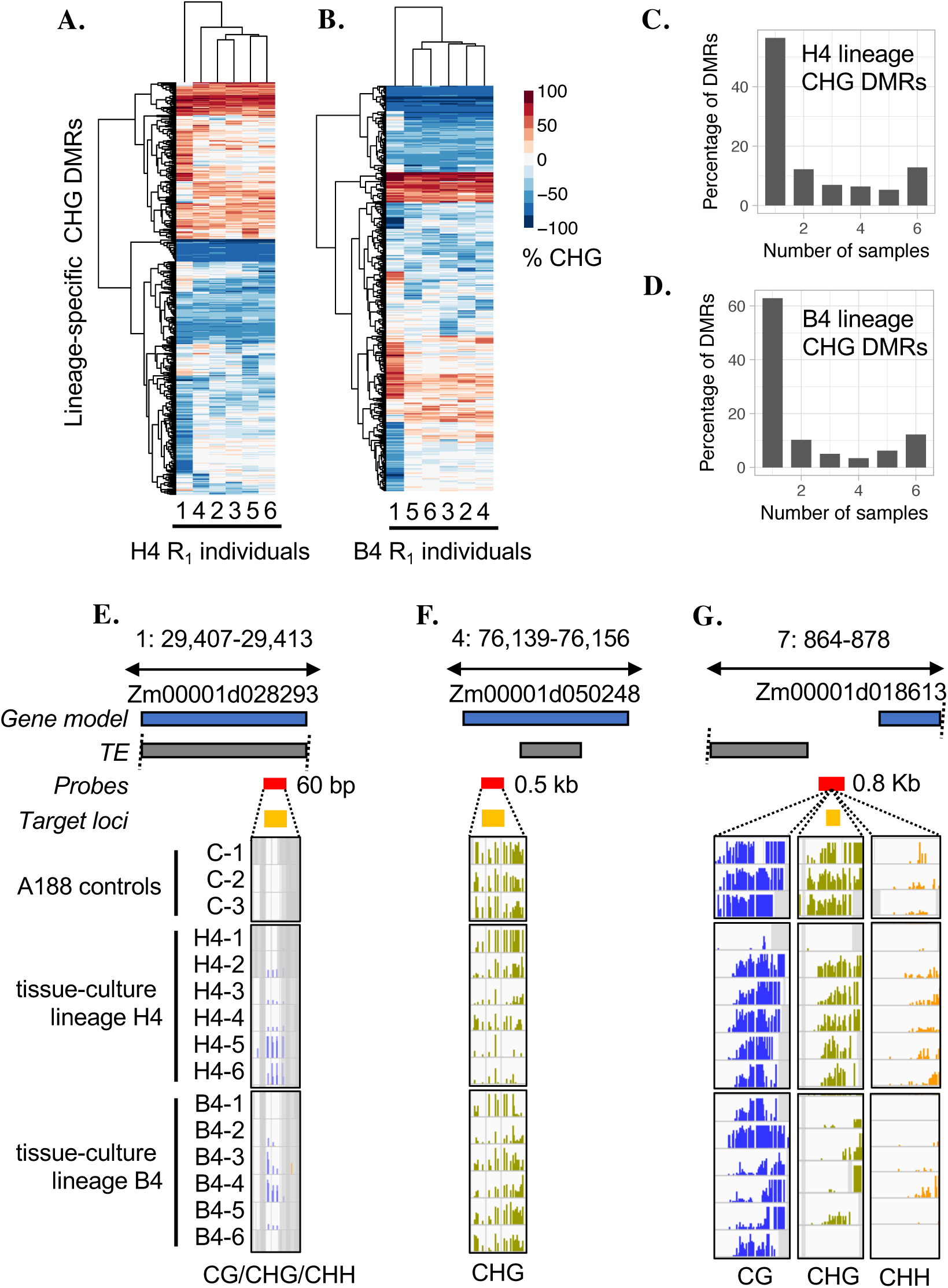
Context specific qualities of DMRs and heritability within lineages (A)-(B) Visualization of the consistency of DMRs within lineage using hierarchical clustering of the change in methylation levels (sample-control) for all DMRs in the CHG context for each lineage (only DMRs for which there was data in all samples were analysed). (C)-(D) The consistency of CHG DMRs in each lineage is shown as a percentage histogram. (E)-(G) Examples of potentially segregating CG-hyper, CHG-hypo and CG-CHG-hypo DMRs. The blue and grey boxes represent the relative positions of gene and TE models (v4 genome annotation); the red and yellow boxes indicate the SeqCap-Epi_v2 capture probes locations and target loci respectively. For each sample track, bar height represents % methylation (0–100%), purple = CG, green = CHG and yellow = CHH. Grey shading indicates missing data not captured.

### Heritability and evidence for targeted remodeling of the methylome

Previous investigations identified variability in the inheritance patterns of DMRs, some of which exhibited stochastic behavior or incomplete penetrance (Stroud *et al*. 2013; Stelpflug *et al*. 2014). This could result from heterozygous changes induced during tissue culture which would segregate in later generations or could result from re-acquisition of the original epigenetic state at one or both alleles following tissue culture. We examined the variation within the CL3 cell lineage and among six sibling R_1_ plants in each of the B4 and H4 lineages to assess conservation or variability of DMRs within related plants and to provide evidence for stochastic behavior or segregation of methylation levels in R_1_ plants.

The DMRs identified in the CL callus sample, it’s R_0_ (CL3) and R_1_ plants (CL3-1 and CL3-2) were compared to document the origin or loss of DMRs throughout this lineage (**Figure S4A-H**). DMRs were classified according to context (CG or CHG), whether they represent gain (hyper) or loss (hypo) of DNA methylation relative to the controls and whether they were found in the majority of samples (consistent vs rare). The majority of the DMRs present in the CL callus sample were detected in the R_0_ and / or the R_1_ plants (**Figure S4A-D)**. Interestingly, there are a number of DMRs that are found only in the R_1_ plants but not in the callus or R_0_ plants. These may represent loci that underwent heterozygous changes during culture that are not detected as DMRs until they become homozygous in the R_1_.

We noted a high level of correspondence for DMRs found among siblings of the B4 and H4 lineages in both the CHG (**Figure 5A-D)** and CG contexts **(Figure S4I-L**). A substantial number of the DMRs within each family are found in at least five of the six individuals. The fact that major changes in methylation are detected (>50%) and that for many DMRs the majority of siblings exhibit these changes suggests a high frequency of homozygous epiallelic change during culture. However, we were interested in assessing whether there was also evidence for either segregation of heterozygous changes during culture or stochastic re-acquisition of DNA methylation. Examples of potentially segregating CG-hyper, CHG-hypo and CG-CHG-hypo DMRs are shown in **Figure 5E-G**. In **Figure 5E** for both lineages there are examples of plants with both substantial and partial gain in methylation. In **Figure 5F**, only the H4 lineage exhibits varying levels of CHG hypomethylation. In **Figure 5G**, there is varying levels of CG and CHG hypomethylation in the B4 lineage; however, here methylation is only partially lost from the left side of the region and could either represent segregation or stochastic reacquisition of DNA methylation seeded by the flanking methylation.

To globally investigate the evidence for either heterozygous changes during culture that could segregate or stochastic re-acquisition of DNA methylation, we assessed the distribution of DNA methylation differences for various sub-groups of DMRs within these families. Consistent DMRs observed in the majority of samples likely reflect homozygous changes. However, in many cases the consistent DMRs might only be observed in 4-5 of the siblings and we investigated whether the other samples might represent a partial change in DNA methylation or whether they had levels similar to the controls **(Figure 6, Figure S5**). The consistent hypomethylated DMRs exhibit a strong peak with all samples showing major loss of DNA methylation, suggesting complete loss and inheritance of DNA methylation in these samples. In contrast, the consistent hypermethylation DMRs have a broader distribution with some samples exhibiting only ~50% gain relative to the controls. This may reflect stochastic loss of the gained DNA methylation at one allele in some plants. Indeed, the examples shown in **Figure 5E-G** may provide examples of stochastic recovery of the original DNA methylation state in some individuals following tissue culture. We proceeded to assess the DNA methylation patterns for all individuals for rare DMRs only observed in one or two members of a family (**Figure 6, Figure S5**). We might expect that these would represent examples in which there is segregation for epigenetic state and the majority of individuals might have intermediate states. We do observe evidence for a broad distribution of methylation levels at these regions. Both hypo- and hypermethylated DMRs exhibit peaks at the extreme values and at 0 (representing no change from control). In addition, there is evidence for a substantial number of plants exhibiting intermediate levels which could reflect heterozygosity. These rare DMRs likely include a subset of changes that are heterozygous in the R_0_ and segregate in the R_1_ generation. These observations suggest that rare DMRs include some examples of changes that were heterozygous in the R_0_ and other examples that represent stochastic changes.

**Figure 6.**
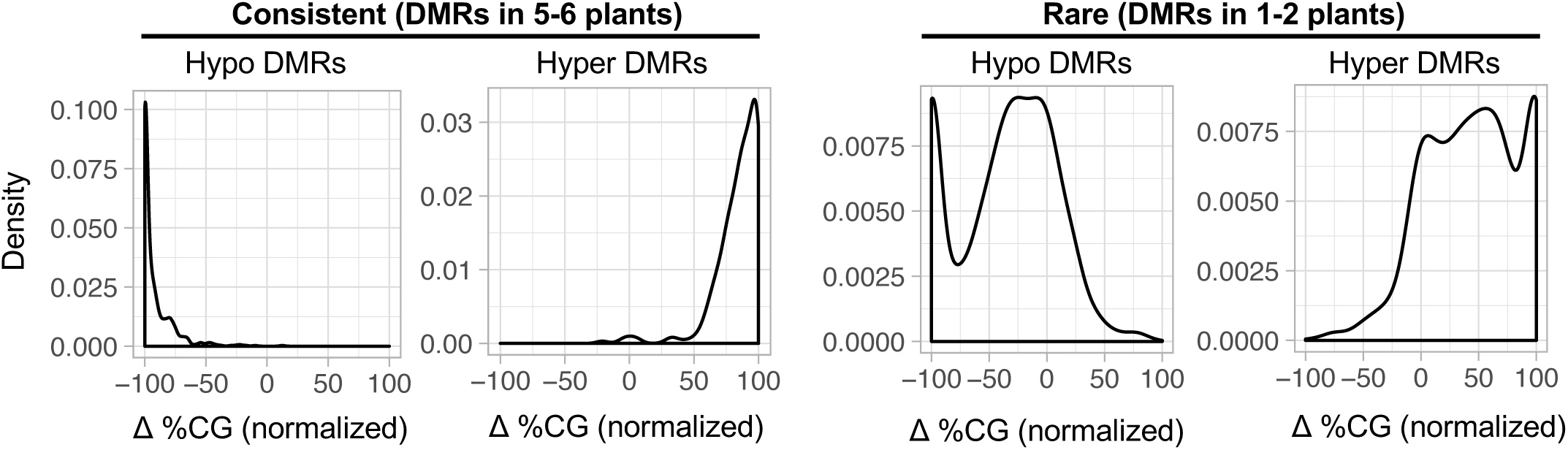
Inheritance and heterozygosity of methylation changes. The distributions of methylation levels for rare and consistent CG DMRs within the H4 tissue culture lineage. Methylation levels were expressed as a differential ratio (Δ %CG) relative to the controls (% CG sample - % CG control), then normalised to the max/min Δ %CG within the lineage by dividing each sample by the value for the sample with the greatest change In methylation (to normalize to 0–100 scale).

## Discussion

DNA methylation can provide additional heritable information beyond DNA sequence in plant genomes. Many studies have found that a substantial proportion of the methylome is highly heritable and not sensitive to environmental perturbation (Song *et al*. 2013; Hagmann *et al*. 2015; Eichten and Springer 2015; Secco *et al*. 2015). However, locus-specific changes in DNA methylation have been observed in response to environmental conditions in several studies (Jiang *et al*. 2014; Wibowo *et al*. 2016). In particular, several studies have found changes in DNA methylation following passage through tissue culture and regeneration (Kaeppler and Phillips 1993; Stroud *et al*. 2013; Stelpflug *et al*. 2014; Ong-Abdullah *et al*. 2015). In this study, we have assessed the context-specific DNA methylation patterns for a portion of the maize genome. In comparison to the significant difference in DNA methylation patterns that can be observed between inbred lines, the plants that had been passed through tissue culture still are quite similar to the control plants of the same genotype. Nevertheless, a principle component analysis reveals evidence for a set of changes that distinguish plants passed through tissue culture.

Tissue culture could impact DNA methylation through a variety of mechanisms. It is possible that tissue culture could simply reduce the efficacy of the DNA methylation, or demethylation, machinery resulting in widespread loss or gain of DNA methylation. While there are more hypomethylation events than hypermethylation events, we do not see major changes in overall DNA methylation levels in plants recovered from tissue culture. Another possibility is that tissue culture simply leads to higher epimutation rates. This would be expected to produce a set of random changes in each event and would also be expected to result in heterozygous changes to DNA methylation. However, we see evidence for significant overlap of DNA methylation changes, even in independent lineages and it seems that the many loci with changes in DNA methylation are affected at both alleles. These observations argue for a targeted process or for specific loci being particularly sensitive to the effects of tissue culture. However, the basis for this targeted mechanism remains unclear. We find both homozygous gains, and losses, of DNA methylation to be more consistent among independent events than expected by chance. In addition, we find that these consistent gains and losses of DNA methylation can affect CHG, CG or both contexts. These observations likely rule out simple hypotheses such as the targeting, or failure to target, specific methyltransferases to certain regions. In addition, we do not find particular features of the target loci that are able to explain why these regions have altered DNA methylation. It is possible that changes in expression at these loci at early stages of tissue culture could alter chromatin and impact the DNA methylation state. Further work will be needed to assess the normal chromatin at these loci and the potential changes during initiation and maintenance of callus.

Thus, a subset of the changes in DNA methylation are consistently found in many independent events and appear to represent homozygous changes in DNA methylation that occur during tissue culture. The majority of these consistent changes in methylation are also observed to occur in natural populations, indicating that the DNA methylation state at these loci is not determined solely by the underlying genetic sequence and suggesting that these are genuine epialleles (Crisp *et al*. 2016). Many natural variation DMRs can be associated with cis sequence variation (Eichten *et al*. 2013) indicative of a mechanism tied to underlying DNA sequence variation. However, given the concurrence of these A188 tissue culture DMRs with natural variants across a wide panel of inbreds, one possible explanation is that this subset of naturally variable loci could be facilitated or pure epialleles in both tissue culture and natural populations (Richards 2006).

The CG-only DMRs often occurred within gene bodies and could reflect the somewhat unstable nature of gene body methylation (Schmitz *et al*. 2013). There is little evidence that changes in gene body CG methylation influence expression levels (Bewick *et al*. 2016; Bewick and Schmitz 2017; Picard and Gehring 2017). In contrast, CHG methylation levels are usually quite stable. The *Karma* locus in oil palm is an example in which a loss of CHG methylation associated with tissue culture results in phenotypic changes (Ong-Abdullah *et al*. 2015). In this case, specific loss of CHG methylation, with no change in CG methylation, at an intronic transposable element results in altered transcripts for the *DEFICIENS* gene and the defective mantled phenotype. Interestingly, we found that many of the CHG-only DMRs identified in this study shared characteristics with the *Karma* locus. These CHG-only DMRs often occurred at transposable elements located within genes. This suggests that the ability to maintain heterochromatin within a largely euchromatin environment may be compromised in tissue culture. There is evidence that changes in CHG methylation without loss of CG methylation can result in changes in gene expression in maize (Anderson *et al*. 2017). Previous studies have suggested that the correct expression and repression of a suite of genes is required for somatic embryogenesis (Salvo *et al*. 2014). It will be interesting to determine whether the inability to maintain CHG methylation within or near genes is a common cause of somaclonal variation in plant species.

## Methods

### Tissue culture and plant materials

All tissue culture experiments were conducted using the regenerable A188 maize inbred. Methylomes of the third seedling leaf of two-week old seedlings from three biological replicates of B73, 2 replicates of Mo17, and single replicates of W22, MoG and Ki11 were captured and analysed for comparison to A188 and regenerated plants. Inbreds, non-cultured control plants and regenerated maize plants of independent maize-immature-embryo-derived cell lines were grown under standard greenhouse conditions at the Gortner Ave. Greenhouse (University of Minnesota, St. Paul, MN). The tissue culture and regeneration process of maize immature embryos was as described by Stelpflug et al. (2014). For methylome analysis, the third leaf from two-week old seedlings was harvested and flash frozen in liquid N_2_ for DNA extraction and sequence capture bisulfite sequencing.

Cell cultures (cell-lines) originated from independent immature embryos named CL, B, D, E, G and H. R_0_ plants were regenerated from each of these cell lines and R_1_ progeny propagated. Each independent R_0_ and R_1_ plant was named using their embryo letter followed by an R_0_ and R_1_ number (eg B4_2 is the second R_1_ progeny of plant 4 regenerated from embryo B). One callus sample named CL, an R_0_ regenerated plant (CL3 - flag leaf), and two R_1_ progeny (CL3_1 and CL3_2) were selected for comparison of different generations. The cell line “CL” was previously described in (Stelpflug *et al*. 2014); however, samples names were modified for consistency. Sample name descriptions herein correspond to the original sample names as follows: callus sample CL = “CL-3”; R_0_ CL3 = “3-7”; R_1_ CL3_1 = “3-7.3”; and R_1_ CL3_2 = “3-7.7”. Seven R_0_ regenerated plants (B4, D1, D2, E6, G7, H3 and H4) from five independent maize cell cultures were selfed, and a total of 25 progeny R_1_ plants were harvested for analysis including 1–6 replicates from each R_0_. In addition, the corresponding 3rd leaf was collected from three different control A188 plants, which were sibling plants originating from the same seed source and were not subjected to tissue culture.

### Maize bisulfite coupled sequence capture (SeqCap-Epi-v2) probe design

The sequence capture probe set was originally designed based on the B73 RefGen_v2 (AGPv2) assembly of the maize genome and subsequently updated to B73 RefGen_v4 aka AGPv4 (also known as Zm-B73-REFERENCE-GRAMENE-4.0) as detailed in **File s1**. A total of 20,643 non-redundant genomic regions spanning 15,728,511 Mb were used to design probes based on the B73 reference genome (AGPv2). These regions were selected based on various criteria. All regions from the first version of capture probes were included in v2 (Li *et al*. 2015a); however the total genome space captured was increased to 15.7 Mb. Additional probes were designed to capture loci satisfying criteria including: DMRs identified between B73, Mo17, Oh43 and between 5 tissues of B73 (Li *et al*. 2015b); tissue culture DMRs (Stelpflug *et al*. 2014); cryptic promoters (Li *et al*. 2015b); mCHH islands (Li *et al*. 2015c); and, various siRNA loci such as phased loci (Zhai *et al*. 2015). The specific target region of interest was termed “specific region” and the bait region captured by the probe design was termed the “target region”. The target regions often included flanking regions of the specific region, and a single target regions can encompass multiple specific regions. A single specific region may satisfy multiple criteria of interest e.g. a region may be both a mCHH island and a CHH DMR. In total, 23,151 “specific” regions (22,950 non-redundant) were defined, including 201 regions that each was annotated to two classes.

### Library construction and sequencing

SeqCap libraries were constructed as described previously (Li *et al*. 2015a). Briefly, DNA of maize-embryo-derived callus, regenerated plants, and non-cultured control plants were isolated using the standard CTAB method. Five hundred to 1,000 nanograms of genomic DNA was used for sequence capture library construction. DNA was sheared to fragments between 180 and 250 bp. The fragments were subject to end repair, dA tailing, ligating to index adapters, and Dual-SPRI Size Selection, followed by bisulfite conversion. The bisulfite converted libraries were then subject to Pre-Capture LM-PCR amplification, purification and quantified using a PicoGreen™ dsDNA Assay Kit. Then the amplified sample library was hybridized to probes designed to target the set of genomic regions selected for analysis. After hybridization, the captured DNA library was bounded to capture beads and the bead-bound DNA was washed. Post-Capture LM-PCR amplification was performed, and following PCR cleanup the libraries were quantitated using PicoGreen. Libraries were pooled together in several batches and the pools sequenced over multiple lanes at the University of Minnesota Genomics Center on a HiSeq2500, in High-output 2×125 bp paired end (PE) mode.

### Data analysis

Adapters were trimmed using *Trim Galore!*. Reads were mapped to maize B73 reference genome AGPv4 using *BSMAP-2.90* (Xi and Li 2009), allowing 5 mismatch or less in a read and quality threshold 20 in trimming 3’end of reads (-v 5 -q 20). Only uniquely mapped reads were kept for subsequent analysis. PCR duplicates were removed using *picard-tools-1.102 ‘*MarkDuplicates’, and *bamtools* was used to remove any improperly paired reads. Overlapping reads were then clipped using ‘bam clipOverlap’ command from *bamUtils*. Conversion rate was determined using the reads mapped to the cytosines of the unmethylated chloroplast genome. The filtered alignment files were then used to derive methylation ratios (i.e., number of methylated and unmethylated reads) for each cytosine in all three sequence contexts (CG, CHG, and CHH) using *BSMAP* tools. The methylation level of a specific target region was calculated based on the cytosines within the region using the weighted DNA methylation method (#C/(#C+#T)). Read coverage per target regions was obtained by counting the number of reads overlapping with the target regions, which was determined using *BEDTools*. Mapping statistics can be found in Table s1.

Following mapping and pre-processing of the raw sequence data, region level methylation data was filtered and processed as follows. Of the 22,749 non-redundant v4 “specific” regions of interest, we obtained data for 21,725 regions in at least 1 sample. Per sample, regions with at least 3 reads were retained. Following 3x read coverage filtering, the mean of the controls per methylation context was calculated for any region that had data for 2 or more control samples. Per context, the number of regions passing the filter were 15,478 (CG), 15,554 (CHG); and 15,751 (CHH); 17,140 unique regions in total. Regions with significant variance among control samples were removed using a strict filter: if control[max - min] was > 30% for CG; > 30 % for CHG and > 10% for CHH. Per context, the number of regions retained after this filter were 13,518 (CG), 13,609 (CHG); and 13,344 (CHH); 15,325 unique regions in total. In total, this yielded 15,325 distinct loci for differential methylation analysis. Per-sample DMRs were identified by comparison to the mean of the control samples where there was a >40% difference in CG/CHG context, and for CHH one sample <5% and one sample >25%. In total, 3921 non-redundant regions had differential methylation in at least one context in at least one sample (**Table s3**). Next the sequence context and overlap of each DMR per sample was determined and DMRs were categorized as CHH, CG-only, CG-CHG and CHG-only. For CG- and CHG-only, a DMR is defined as context specific (“-only”) where the change in the other context is less than 10% compared to the controls (eg “CG-only hypo” occurs where CHG does not drop by more than 10%).

Lastly, all CG and CHG DMRs were aggregated into a non-redundant list and consensus calls across all samples were evaluated as described in **Table s3**. A consensus direction for each region was determined if there was a greater than 80% consensus among samples regarding the direction of the DMR call. The consistency of each DMR was then determined by calculating the percent support for each DMR among the samples. Finally, a specificity consensus of the DMR was determined, indicating whether the DMR is specific to a single context and occurring without a significant change in the other context. PCA analysis was performed in R using the *pcaMethods* package, using data summarized to 100 bp tiles for each sample. Gene feature files (GFF) were downloaded from MaizeGDB and the Transposable Element annotation file from (Jiao *et al*. 2017).

### Data Availability

**File S1** contains detailed descriptions of the conversion between B73 RefGen_v2 assembly of the maize genome and B73 RefGen_v4. The annotation of the sequence capture design of SeqCap-Epi-v2 is available under the DOI: https://doi.org/10.13020/D69X0H. Sequence data will be uploaded to the NCBI Short Read Archive and BioProject database.

**Table S1**

Sample metadata and sequencing metrics. Capture batch refers to the in house batch code; conversion rate calculated using chloroplast mapping reads; On target metrics calculated using Picard tools.

**Table S2**

Identification of context-specific DMRs. For CG and CHG methylation, DMRs are defined as regions where the average DNA methylation percentage levels differ by greater than 40% in a sample relative to the control average. DMRs in the CHH context were identified when one sample has >25% and another has less than 5% CHH methylation.

**Table S3**

Non-redundant list of CG and CHG DMRs. Columns B-J represent the aggregation of CG and CHG DMR calls per sample. Column K-P data for calling consistency of DMRs; “samples_3_reads” is the total number of samples with sufficient data (3 reads) to make a DMR call for that region; “direction_consensus” refers to whether there was a greater than 80% consensus among samples regarding the direction of the DMR call for hypo, hypo or else a bidirectional; “DMR_calls” is the total number of samples in which the region was identified as a DMR; “support %” = “DMR_calls”/”samples_3_reads”*100. If there was >= 50% support for a DMR between samples it was classified as “consistent”, all others were considered “rare”. The columns Q-U are data for calling specificity consensus of the DMR, indicating whether the DMR is context specific and occurring without a significant change in the other context. A DMR is defined as context specific where the change in the other context is less than 10% compared to the controls (see Figure X). Q-T represent the aggregation of per sample specificity calls; “specificity” is the consensus call if >= 80% of the call agree.

**Table S4**

Gene loci that may lose heterochromatin. our sequence capture design profiled methylation information at 1,447 gene loci that have high CG and CHG methylation in the control A188 plants indicative of heterochromatin (control average >50% CG and >50% CHG). We detected 219 CHG-hypo DMRs that overlapped these loci, corresponding to 200 distinct genes. Functional annotations were downloaded from Gramene release B73_v4.

## Acknowledgments

We thank Peter Hermanson for sequence-capture library preparation and technical support; Yaniv Brandvain for statistical advice; and, Jackie Noshay, Sarah Anderson and Peng Zhou for legendary computational assistance. This work was funded by a grant from the National Science Foundation (IOS-1237931) to NMS. Illumina sequencing was performed at the University of Minnesota Genomics Center. Computational support was provided by the Minnesota Supercomputing Institute and the Texas Advanced Computing Center.

---

**Figure S1.**
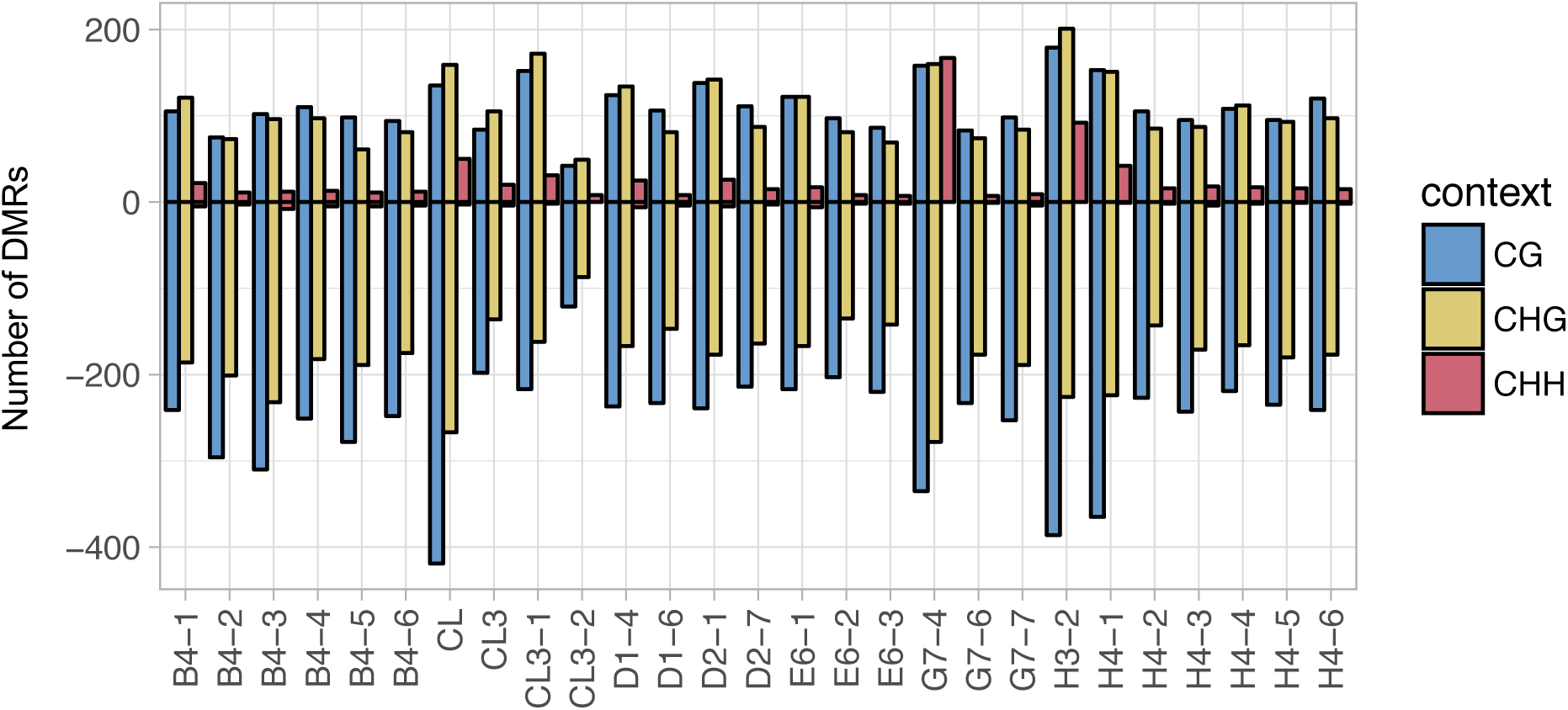
Identification of altered methylation in CG, CHG and CHH contexts following tissue culture. The total number of DMRs per context identified for each sample passed through tissue culture. Bars indicate the total number of DMRs for both hypermethylated (positive bar) and hypomethylated (negative bar) in each sample relative to A188 control plants. For each sample DMRs were identified by comparing to the average of the three controls samples (after removing stochastic regions among controls) to each tissue culture sample for those regions with a minimum 3 mapping reads. DMRs defined as a methylation percentage difference of 40% or greater for CG and CHG, and for CHH a 25% or greater difference and one sample less than 5% methylation.

**Figure S2.**
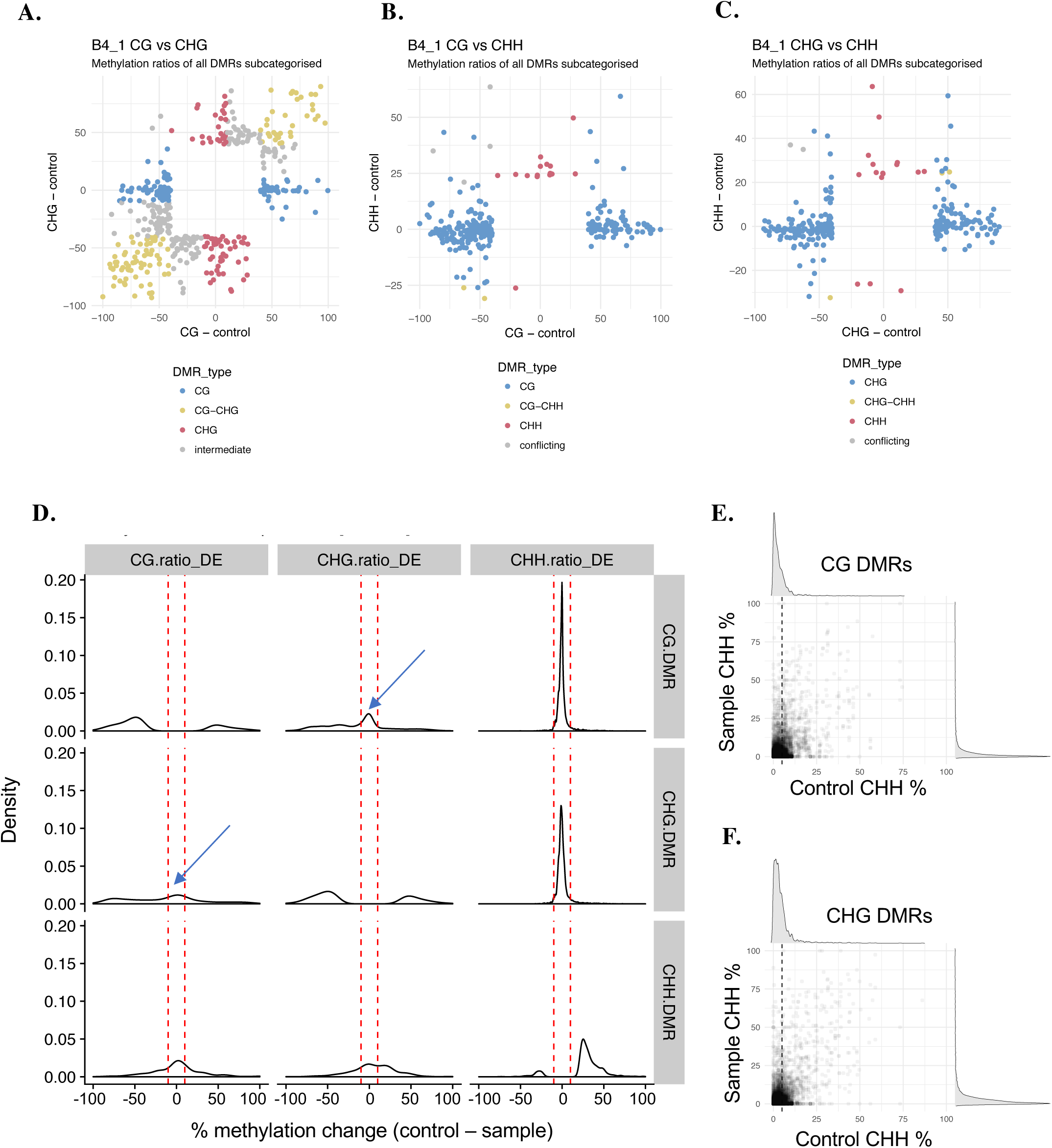
Identification of context specific DMRs. (A)-(C) Overlap of CG, CHG, CHH DMRs for the individual sample, B4–1. Colours indicate DMR class, values represent percent methylation change calculated as (% methylation control) - (% methylation sample). (D) Density plots of methylation ratios (sample – control) for tissue culture sample B4–1 for regions identified as DMRs in a particular context. This plot illustrates that for CG and CHG DMRs the level of CHH methylation changes very little. CG and CHG levels exhibit two behaviours: they can change independently (context specific DMRs) as indicated arrows for the areas between the red lines; and also concurrently. (E)-(F) Percent CHH methylation at each CG and CHG DMRs is plotted in the controls compared to the samples. There are very few examples of changes in CHH methylation levels at CG or CHG DMRs and in most cases the level of CHH methylation at these regions is very low in both the control and tissue culture samples.

**Figure S3.**
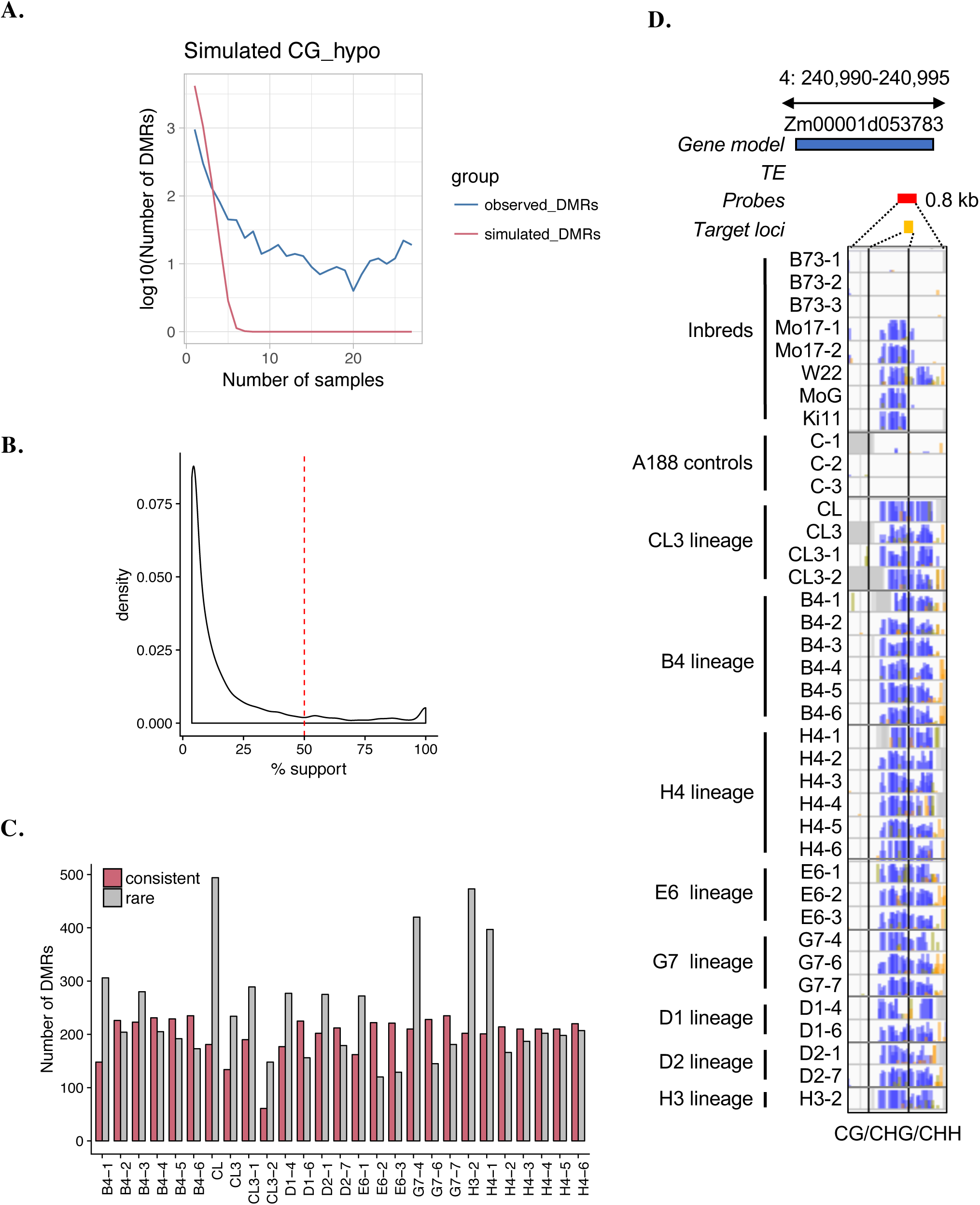
Methylation changes induced by tissue culture are common to many independent samples. (A) A simulation was performed to document the expected frequency of DMRs occurring in multiple samples (red line). In contrast, the observed DMRs are enriched for occurrence in multiple samples (blue line). To simulate the expected overlap of DMRs between different plants by random chance, regions were randomly selected, per sample (to reflect the total number of DMRs and regions profiled per sample), and the random overlap of DMRs is plotted in the red line above. The simulation was replicated 100 times and averaged. The blue represents the actual observed number of recurring DMRs between the tissue culture samples. (B) The percent of samples (with data) supporting each DMR was determined. We divided the DMRs as consistent (occurring in >50% of samples) or rare (occurring in <50% of samples). (C) Total number of rare (grey bars) and consistent (red bars) DMRs observed per sample. (D) Example consistent CG DMR observed in all tissue culture samples in the gene body of Zm00001d053783. This DMR has a low methylation level in A188 and B73 but is higher in other inbreds and in all samples that have experienced tissue culture. The blue and grey boxes represent the relative positions of gene and TE models (v4 genome annotation); the red and yellow boxes indicate the SeqCap-Epi_v2 capture probes locations and target loci respectively. For each sample track, bar height represents *%* methylation (0–100%), purple = CG, green = CHG and yellow = CHH. Grey shading indicates missing data not captured.

**Figure S4.**
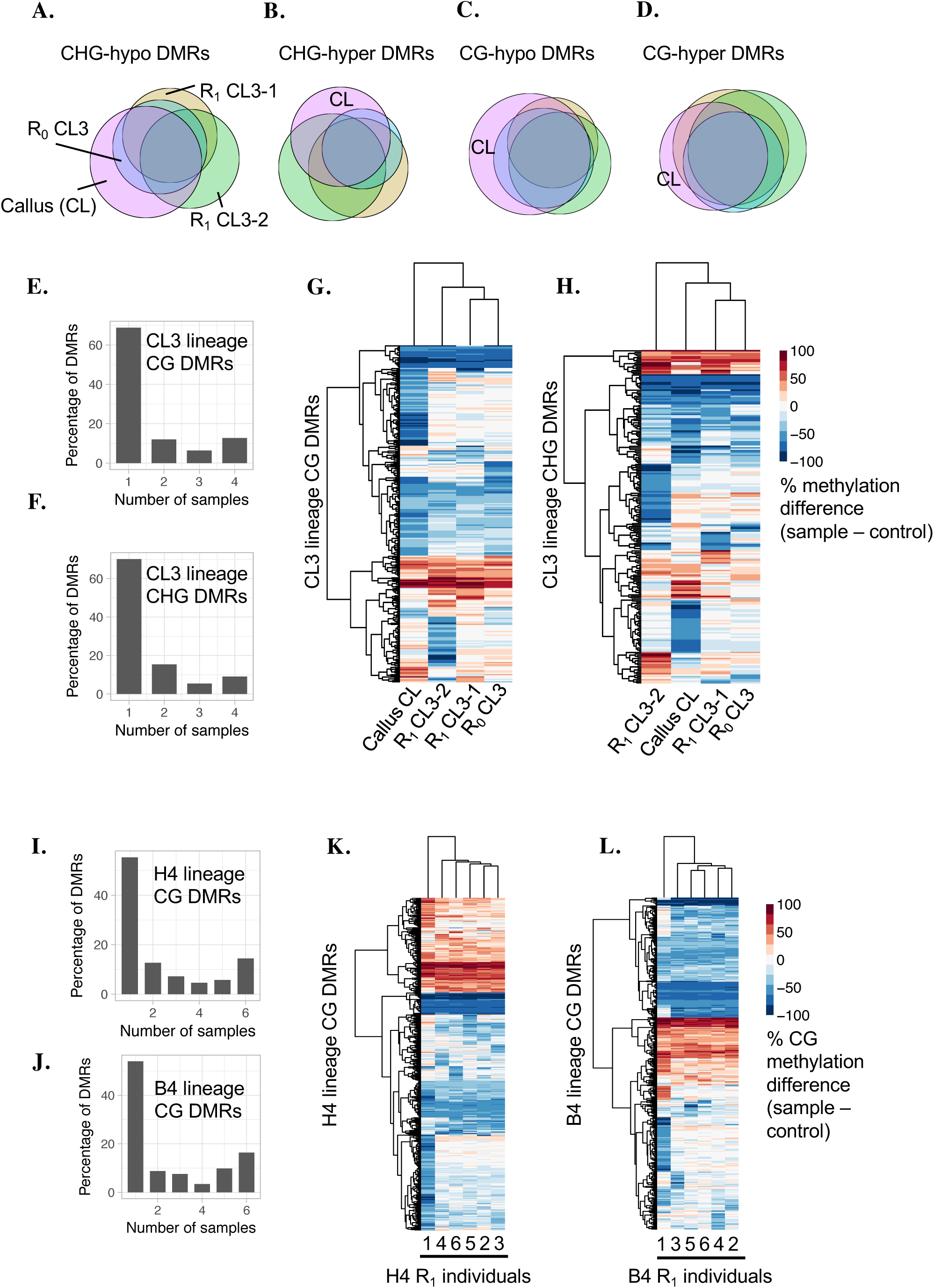
Context specific qualities of DMRs and heritability within lineages. (A)-(D) Overlap of DMRs among plants in the CL3 lineage represented as proportional Venn diagrams. DMRs are divided into CG/CHG and hypo/hyper; the colour corresponding to each sample is indicated in (A). (E)-(H) Analysis of the CL3 lineage; DMR consistency histograms displaying the distribution of the number of samples for; (E) CG and (F) CHG DMRs identified in for the CL lineage. (G)-(H) Hierarchical clustering of the change in methylation levels (sample - control) for all DMRs in the CG and CHG contexts for the CL lineage. (I)-(L) Analysis of the H4 and B4 lineages; DMR consistency histograms displaying the distribution of the number of samples for; (I) H4 CG and (J) B4 CG DMRs. (K)-(L) Hierarchical clustering of the change in methylation levels (sample - control) for all DMRs in the CG contexts for the H4 and B4 lineages. For each analysis, only DMRs for which there was data for all samples were considered.

**Figure S5.**
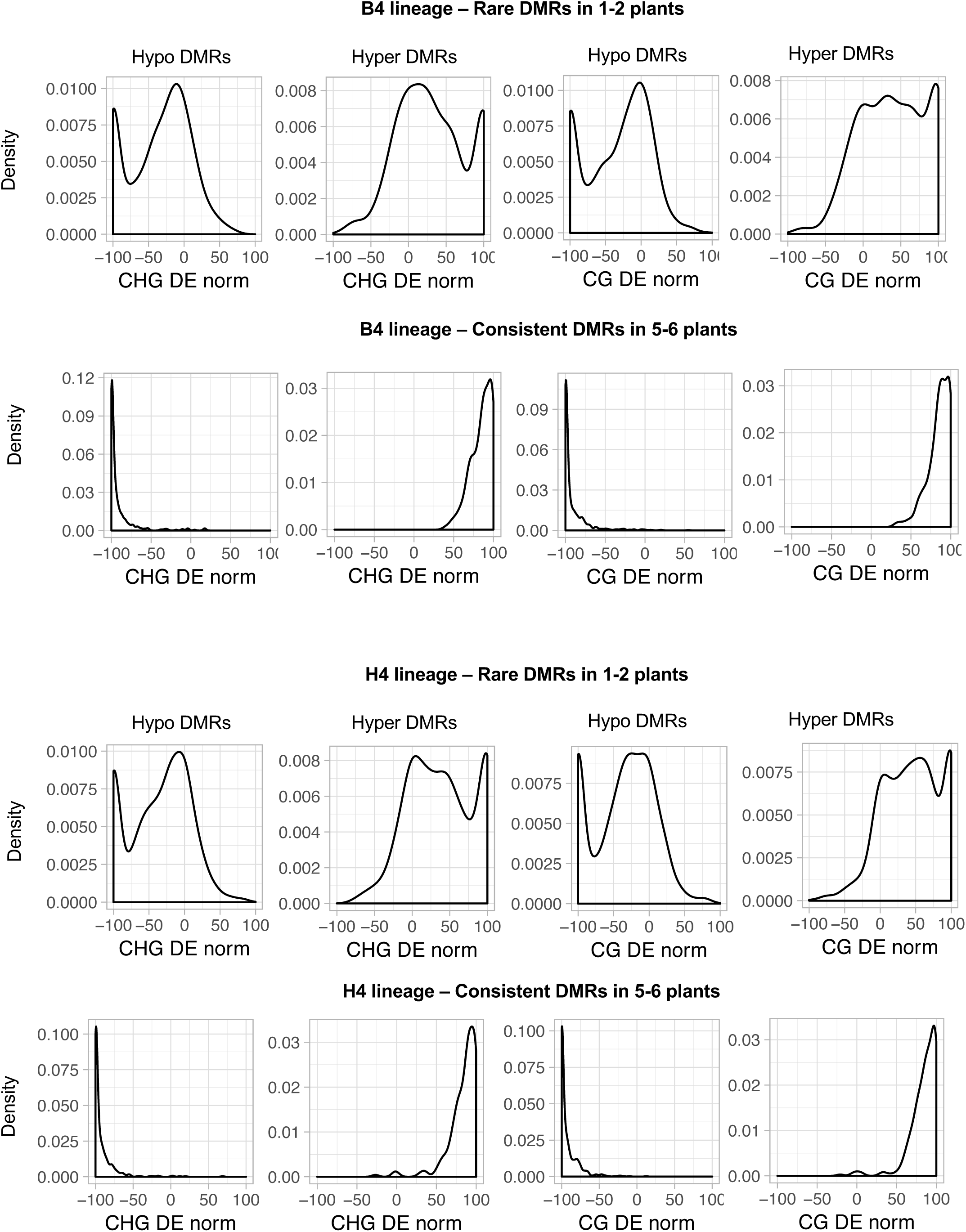
Inheritance and heterozygosity of methylation changes. To investigate whether there is a heterozygosity signal in the methylation data, the distributions of methylation levels for CG and CHG DMRs within tissue culture lineages were examined. Methylation levels were first expressed as a differential ratio (DE mC) relative to the controls, then normalised to the max DE mC within the lineage (DE norm). Only DMRs for which there was data for all samples were considered. If all sites were homozygous in the callus/R_0_ then there would be a single peak at 100 in the R_1_ data; whereas if all sights were heterozygous in the R_0_ we would expect three peaks at 0, 50 and 100 in a ratio 1:2:1. The data is likely a mix of both signals and further diluted by partial loss or gain of methylation over time.

